# Combining video telemetry and wearable MEG for naturalistic imaging

**DOI:** 10.1101/2023.08.01.551482

**Authors:** George C. O’Neill, Robert A. Seymour, Stephanie Mellor, Nicholas Alexander, Tim M. Tierney, Léa Bernachot, Mansoureh Fahimi Hnazaee, Meaghan E. Spedden, Ryan C. Timms, Daniel Bush, Sven Bestmann, Matthew J. Brookes, Gareth R. Barnes

**Author notes:** Corresponding author: *George C. O’Neill:.

## Abstract

Neuroimaging studies have typically relied on rigorously controlled experimental paradigms to probe cognition, in which movement is restricted, primitive, an afterthought or merely used to indicate a subject’s choice. Whilst powerful, these paradigms do not often resemble how we behave in everyday life, so a new generation of ecologically valid experiments are being developed. Magnetoencephalography (MEG) measures neural activity by sensing extracranial magnetic fields. It has recently been transformed from a large, static imaging modality to a wearable method where participants can move freely. This makes wearable MEG systems a prime candidate for naturalistic experiments going forward. However, these experiments will also require novel methods to capture and integrate information about complex behaviour executed during neuroimaging, and it is not yet clear how this could be achieved.

Here we use video recordings of multi-limb dance moves, processed with open-source machine learning methods, to automatically identify analysis time windows of interest in concurrent wearable MEG data. In a first step, we compare a traditional, block-designed analysis of limb movements, where the times of interest are based on stimulus presentation, to an analysis pipeline based on hidden Markov model states derived from the video telemetry. Next, we show that it is possible to identify discrete modes of neuronal activity related to specific limbs and body posture by processing the participants’ choreographed movement in a dancing paradigm. This demonstrates the potential of combing video telemetry with mobile neuroimaging for future studies of complex and naturalistic behaviours.

## 1) Introduction

One of the challenges for neuroimaging is the development of more *naturalistic* and ecologically valid approaches for studying brain function (Finn et al., 2022). Much effort has been made in creating complex visual stimuli to assess brain function in a way which is closer to the real world than the laboratory (Betti et al., 2013; Kim et al., 2018; Kringelbach et al., 2023). However, a fundamental limitation remains in that the actual expression of behaviour through movement is limited in most neuroimaging modalities. This, ultimately, limits the ecological validity of studies of brain function.

Magnetoencephalography (MEG; Cohen, 1972; Hamalainen et al., 1993) is a powerful non-invasive method of imaging neural function from the brain, which measures the changes in extracranial magnetic field originating from intra-cellular current flow. As MEG is a direct measure of underlying neural processes, it offers a millisecond scale temporal resolution to capture the rapid dynamics of cognition. Unlike electroencephalography (EEG) which measures the same signal via electric potentials on the scalp, magnetic fields pass through the skull with relatively little distortion, allowing MEG to offer a higher spatial resolution when the data are source-reconstructed. However, magnetic fields from neural populations in the brain are of the order of 10s of femtoTesla, so traditional MEG systems have required an array of highly sensitive superconducting sensors and cumbersome cryogenic infrastructure to support them. This means that participants have to keep their head (and by extension, the rest of their body) as still as possible within a gantry which contains the sensing array. Whilst some attempts at naturalistic MEG studies have been made (e.g. listening to speech or watching movies; Betti et al., 2013; Nunes et al., 2020; Park et al., 2016; Thiede et al., 2020), the constraint on movement makes these studies passive, with participants not behaving in a natural way during the presentation of the stimulus.

Recent advances in the engineering of cryogen-free sensors, particularly optically pumped magnetometers (OPMs; Knappe et al., 2016; Osborne et al., 2018; Shah and Wakai, 2013; Sheng et al., 2017) have led to the rapid development of a new generation of optically pumped (OP) MEG systems (Borna et al., 2017; Boto et al., 2017; livanainen et al., 2019; Pratt et al., 2021). OP-MEG has been used to assess sensory (Hill et al., 2019; Rier et al., 2024; Roberts et al., 2019; Seymour et al., 2021) and cognitive processes (Barry et al., 2019; de Lange et al., 2021; Rhodes et al., 2023; Tierney et al., 2018), as well as showing its potential for clinical applications (Feys et al., 2023, 2022; Hillebrand et al., 2023; Vivekananda et al., 2020). The diminutive size of commercial OPMs (approximately the size of a 2-by-3 LEGO brick) mean that many of these sensors can be packed into a rigid helmet (Boto et al., 2017) or soft cap (Feys et al., 2022; Hill et al., 2020), close to the scalp and worn in a form factor similar to EEG. A further advantage is the fact that muscle activity from the neck or other body parts do not contaminate the MEG signal to the same proportion as in EEG (Boto et al., 2019; Muthukumaraswamy, 2013), which can prove highly beneficial if a study requires explicit movement. Indeed, OP-MEG has already demonstrated compatibility with immersive virtual reality environments (Roberts et al., 2019) and used to reliably image neural processes whilst participants move large (> 1 m) distances (Holmes et al., 2023b; Mellor et al., 2023; Seymour et al., 2021), play a ball-game against each other (Holmes et al., 2023a), or simply drink tea (Boto et al., 2018). It is for these reasons that OP-MEG promises to be a powerful tool in the field of naturalistic imaging with unconstrained movement (Stangl et al., 2023).

One challenge to this approach, however, is how to incorporate complex behaviour into the modelling of neuroimaging data as experiments become increasingly realistic (Stangl et al., 2023). Recent developments in computer vision and machine learning methods to track behaviour offer a promising approach (Anderson and Perona, 2014; Bigand et al., 2024; Kaneko et al., 2024; Mathis et al., 2018; Schneider et al., 2023; Weinreb et al., 2023). Here, we demonstrate how behavioural information that is extracted from data-mined video of participants performing complex motor paradigms can be fused with concurrently recorded wearable MEG data to yield brain activity maps of complex behaviour. We first show that we can decode the experimental state of the participant from the video data alone. Next, we show that by using the video, we can return context to MEG data from participants dancing and use the video-derived cues to quantify neuronal activity associated with specific limbs or postures. In sum, we describe a largely automated method of processing video data to identify time windows of interest in concurrently recorded neuroimaging data, thereby providing an analysis pipeline to support future naturalistic experiments across a range of cognitive domains

## 2) Methods

### 2.1) Experiment

This study was carried out at the OP-MEG scanning suite at the Wellcome Centre for Human Neuroimaging. The project was approved by the University College London research ethics committee. All participants who took part in the study provided informed written consent prior to MEG/MRI data collection.

#### Motor block design

Three participants (all male, aged 43±12 [mean±SD years]) took part in a motor paradigm. Seated in the middle of the magnetic shielded room (MSR), the participant was visually cued to move one of their four limbs freely until a fixation cross appeared on the screen. The movement epochs were 4 s in length with a 10-11 s inter trial interval. In a block, 15 trials of each condition were presented in a pseudorandom order, with two blocks recorded per participant.

#### Dancing

Five participants (all male, aged 37±12 [mean±SD years]) danced the *Hokey Cokey*, a popular dance for school children in the United Kingdom. It was selected as its major actions involve moving individual limbs separately, with periods where the lyrics explicitly instruct the dancer how to move (“you put your left arm in…”) as well as periods of ambiguity (“whoa, the Hokey Cokey!”). Participants were given basic choreography training prior to recording. An audio recording of the song (Black Lace, 1985) was played into the room from a set of speakers placed outside of the room. Participants repeated the dance multiple times. In total 20 recorded dances across all participants were kept for further analysis.

### 2.2) Acquisition

#### Magnetic Resonance Imaging

Each participant underwent Magnetic Resonance Imaging (MRI) in preparation for the study on a Tim Trio 3T MR System (Siemens Healthineers, Erlangen, Germany). Two images were acquired for each participant. The first, a modified FLASH sequence with a high-bandwidth readout (FOV: 256 mm (A-P) x 256 mm (S-I) x 192 mm (L-R); resolution 1 mm × 1 mm × 1 mm) for minimal distortion of the participants’ face and scalp, whilst maintaining enough dynamic range to segment white and grey matter in the brain. Full details of the acquisition parameters can be found in Meyer et al., (2017). The second, a T1-weighted image (MPRAGE; TR=2530 ms, TE=3.34 ms; FOV: 256 mm (A-P) x 256 mm (S-I) x 172 mm (L-R); resolution 1 mm × 1 mm × 1 mm) was collected to supplement the first scan, in case automatic segmentation methods with the FLASH MR image failed.

#### MEG

The OPM arrays consisted of a combination of 2^nd^ generation (dual axis) and 3^rd^ generation (triaxial) zero-field magnetometers (GluSpin, Louisville, CO). The dual axis sensors provided axial-oriented field detection and one tangential field measure, the triaxials give a full vector field measurement. The sensors operated in an open-loop mode with an operational dynamic range of ∼±4.5 nT relative to their zero-field point. The number of channels recorded ranged from 64 to 128; detailed breakdowns of channel counts and sensor layouts in the Supplementary Material. The MEG data were digitised using a 16-bit ADC system (National Instruments, Austin, TX) at a sample rate of 6Q00 Hz.

The sensors were placed in bespoke 3D printed scanner-casts (Chalk Design, London, UK) specifically designed for each participant (Boto et al., 2017; Tierney et al., 2018). Scanner-casts ensure a comfortable fit and minimise the coregistration errors between the sensors and the participant’s anatomy (Meyer et al., 2017) The helmet’s geometry was based on a scalp mesh extracted from the FLASH MR images. The OPMs were oriented such that the manufacturer-defined Y-axis measured the component of the magnetic field axial/radial to the scalp, with the centre of the vapour cell typically between 9 and 12 mm from the scalp surface. Sensor locations were in the same coordinate system as the anatomical images due to the manufacture process of the scanner-casts, so no additional registration was required.

MEG data were acquired in an MSR (Magnetic Shields Ltd, Staplehurst, UK), with internal dimensions of 4380 mm × 3380 mm × 2180 mm. The MSR is constructed from two inner layers of 1 mm mu-metal, a 6 mm copper layer, and then two external layers of 1.5 mm mu-metal. The room contains a series of built-in degaussing coils to minimise the residual background field in the room (Altarev et al., 2015). The degausser was used in the period after closing the participant in the room and prior to data acquisition. No external active magnetic shielding was used for these experiments.

#### Video

Visible-light spectrum video recordings of the participants performing the experiments were recorded alongside the MEG data using a camera attached to a single-board computer (Raspberry Pi Foundation, Cambridge, UK). The camera was triggered to record via a GPIO pin controlled by the stimulus presentation software, sent to both the camera and the OPM acquisition electronics to allow for offline synchronisation. Video was recorded at a resolution of 640 × 480 pixels with a frame rate of 30 Hz (with the exception of subjects 004 and 005, which were recorded at 50 Hz). The camera was located in front of participants 001, 002 and 005 (see Fig. 1 for an example), and approximately 45 degrees off axis to the right of the participants 003 and 004.

**Figure 1.**
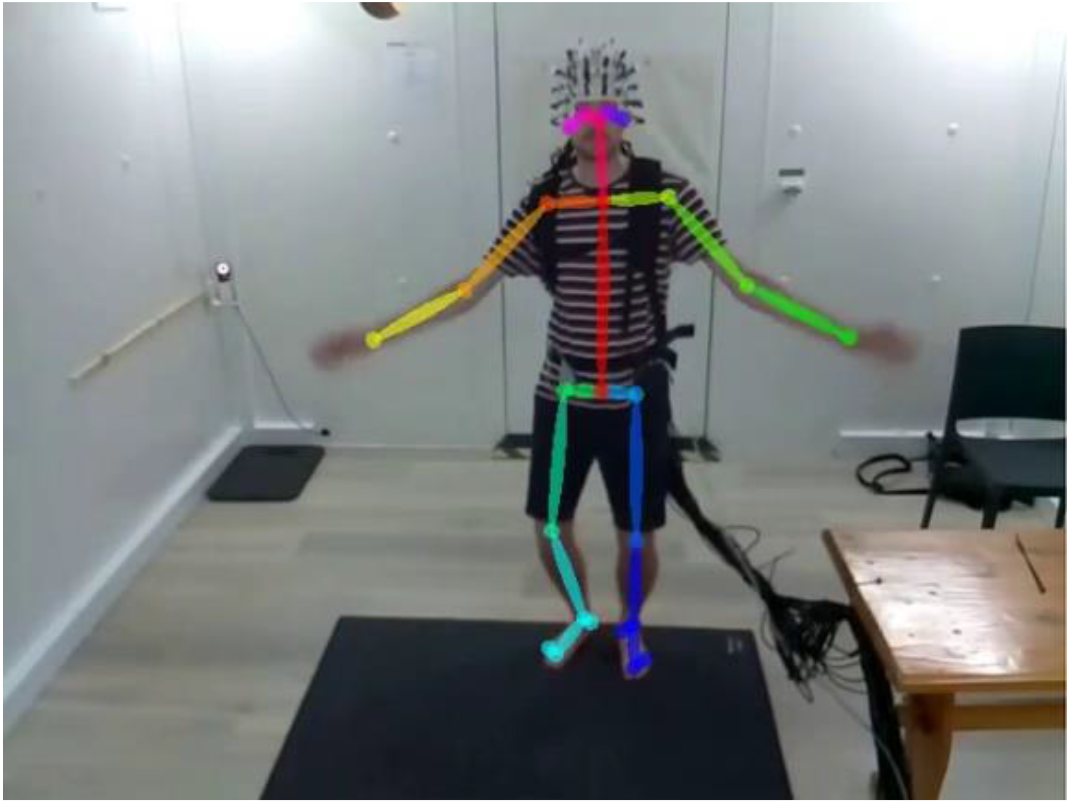
A subject performing the dancing paradigm within the magnetically shield room whilst wearing the MEG system. The pose estimation results for this video frame have been superimposed to represent where the keypoints (circles) used for telemetry analysis are located.

### 2.3) Preprocessing

#### Video

Telemetry information from the video was extracted using OpenPose (Cao et al., 2021), a pre-trained convolutional neural network which is capable of single and multi-person pose estimation from images and video. Inference was performed using an RTX A5000 GPU (NVIDIA, Santa Clara, CA), and for every frame, the X-Y pixel locations and a confidence-of-fit score for 25 different key-points of the body were recorded. An example of the keypoints fitted to each frame can be found in Figure 1. For each experiment, the telemetry position data was linear interpolated to fix missing data, was converted to speed (units: pixels *I* frame) and normalised via a Z-transform. Finally, the 50 Hz telemetry from participants 004 and 005, were linear interpolated to match the 30 Hz video sampling rate of subjects 001-003. For the dancing telemetry, each session’s data were normalised individually and then concatenated in time. The data were then partitioned using a hidden Markov model (HMM), with a multivariate Gaussian mixture model used as the observation model, to derive when differing movement ‘states’ occurred in the data, HMM inference was handled with the HMM-MAR toolbox (Vidaurre et al., 2016) to separate the data into *k* states. On completion, the fitted HMM returned a probabilistic timeseries for each of the *k* states and Viterbi path which assigns a mutually-exclusive state on a per-frame basis.

In the block design experiment, we set *k =* 15. To identify what each state corresponded to, the binarised Viterbi path for a given state was compared to the initial block design timings using the Jaccard index. The Jaccard index is a measure of the intersection of two sets relative to their union (Jaccard, 1912). The state with the highest Jaccard index for a given experimental condition was labelled with that condition name. The 11 remaining (unlabelled) states were considered as rest and combined into a single rest *meta-state*.

For the dancing experiment, where the video consisted a concatenated group of subjects (from different angles in the room) we fitted a larger, k=25 state HMM. The higher number of requested states would allow for multiple states corresponding to the same limb (but from different subjects) to be identified. After HMM fitting, the Viterbi paths were regressed against the telemetry of each key-point, to generate regression maps of movement (examples can be seen in Figures 2B and 4A). From these maps, which limb (or dance move) a state corresponded to was performed by visual inspection, with appropriate labelling of the states applied. If two or more states represented the same limb (but across different subjects), their Viterbi paths were pooled together to make one meta-state representing the limb across all subjects (details of state -limb mappings can be found in the Supplementary Material).

**Figure 2.**
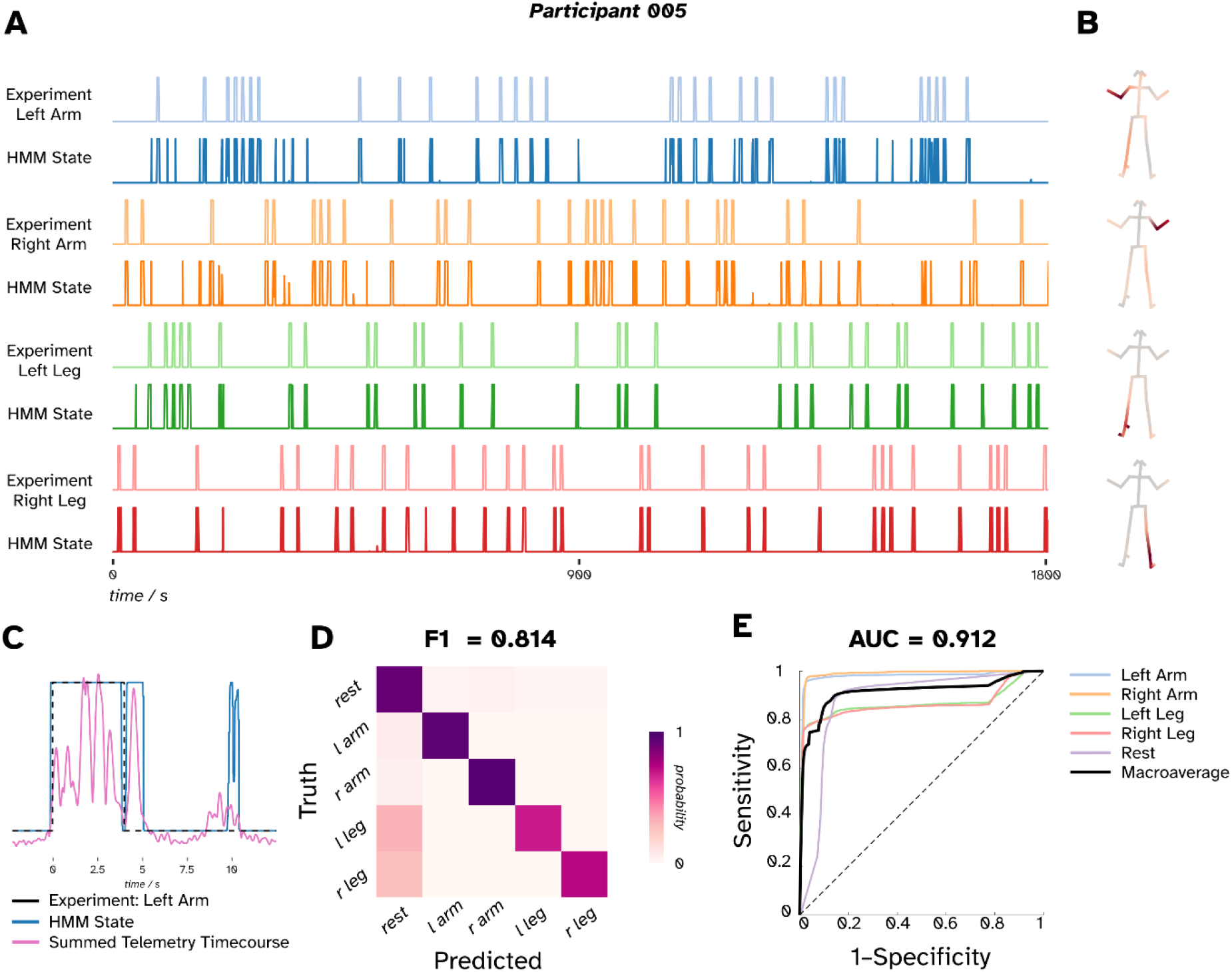
A comparison of the HMM-derived video telemetry states in comparison to the original experiment design for a simple 4 condition motor task in a single subject (participant 005). **A**) Temporal plots depicting the 4 experimentally derived states across a recording represented as binary signals. Plotted below each experimental timecourse is the corresponding telemetry state which closest matched each experimental condition. The duration of each movement was 4.2 ± 0.9 (mean ± SD) s. **B)** Heatmaps corresponding to the regression of the HMM state timecourse against the original telemetry data to identify which body parts were moving in a given state; the darker the plot, the stronger the relationship between telemetry data and a given HMM state. **C)** A portion of the left arm experimental state (black dashed line) and the closest matched HMM state (blue solid line) which demonstrates the delay (∼1000ms offset) between presentation of experimental cue and cessation of movement, l/l/e also see a period of small movements (pink line) occurring at around 8 seconds (during the inter-trial interval) which classified as movement by the HMM, D) Multi-class confusion matrices showing the probability of a given HMM state being active compared to the experimental ground truth. E) Receiver operating characteristic (ROC) curves for the resultant HMM compared to the block design it is predicting. Pale colours represent the states representing specific conditions in a (one v. all) classification, black line represents the macroaverage (unweighted mean) of all 5 ROC curves.

#### MEG

An anti-aliasing 500 Hz low-pass filter (60^th^ order FIR filter combined with a Kaiser window) was applied to the MEG data and then downsampled to a sample rate of 2000 Hz. The HMM state time courses derived from the video were synchronised to the MEG using nearest neighbour interpolation and appended to the dataset as supplementary trigger channels. Environmental interference was reduced by applying a Homogenous Field Correction (HFC), where the interference is modelled as a set of regular spherical harmonics derived from the sensor array (Tierney et al., 2022). Here, 8 components were projected out of the data (3 homogenous field components and 5 linear gradients). If any channels had a residual variance larger than WQ pT after HFC, the projection was undone, these channels were disabled and the HFC was reapplied again without those channels included in the model. The sensor compensation matrix was also updated to account for the linear combinations of sensors in the forward modelling (Tierney et al., 2021). A secondary advantage of using HFC is that its performance is robust across arrays of differing channel numbers and types. The expected differences in signal is <1dB across the arrays utilised (Tierney et al., 2022).

### 2.4) Source localisation

Source localisation was performed with the DAiSS toolbox supplied with SPM12 (Litvak et al., 2011). Sources were modelled along a 5 mm grid within a boundary delineating the brain and CSF of the participant. The forward model was Nolte’s single shell (Nolte, 2003), where the conductive volume geometry was the same brain/CSF boundary. Three dipoles per location representing the cardinal orientations were generated, with their degrees of freedom reduced to two to compensate for the ‘silent radial source’. A linear-constrained-minimum-variance (LCMV) beamformer (Brookes et al., 2008; Van Veen et al., 1997) was implemented to perform the inverse modelling. Here, the data covariance matrix was generated with the MEG recording band-pass-filtered into the 830 Hz band, which has been strongly associated with sensorimotor activity (Gaetz et al., 2020; Jurkiewicz et al., 2006; Pfurtscheller and Lopes da Silva, 1999). Due to the projection carried out during HFC in preprocessing, the MEG data are rank deficient. To control for this, the covariance matrix was regularised; the matrix decomposed into a set of eigenvectors and eigenvalues, with the 8 smallest eigenvectors/eigenvalues (associated with the 8 projected-out components from HFC) discarded (Westner et al., 2022). During LCMV, the forward models for each source were linearly combined to maximise variance from that location. (Sekihara et al., 2004).

### 2.5) First-level source analysis

To map which sources in the brain covaried with the onset and offset of the behavioural states, we opted to take a 2-level general linear model (GLM) approach (Brookes et al., 2004; Quinn et al., 2024; Worsley and Friston, 1995). In particular, we took an approach based on the GLM-beamfomer (Brookes et al., 2004). For the first level, we performed a mass-univariate multiple regression on each source. For each source we band-pass filtered the reconstructed source timeseries between 8-30 Hz and generated an amplitude envelope timecourse via a Hilbert transform. Prior to regression, the envelopes were log-transformed (Hawkins and Wixley, 1986). A given processed source signal 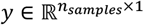 is fitted to our general linear model:

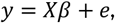

where, 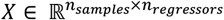, is our design matrix with our temporal features of interest in the columns, 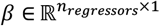 are the regression coefficients, and 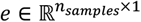 is our unexplained data. How we constructed *X* for each experiment differed as follows. For the motor experiment, we generated three design matrices, where the first column was either one of three features of interest. These features of interest were: the active period as specified by the block design (box car between 0-4 seconds of trial onset), a baseline ‘rest’ period (box car between 7-11 seconds of trial onset) or the HMM-derived active period (binarised state time-series). The second column modelled was the mean. A regression was performed for each source in the brain, generating a set of regression coefficients per vertex, which were combined to generate images. For each trial, three regression coefficient images (block active, block rest, and HMM-active) were kept for further analysis.

For the dancing experiment design, as our motor states were orthogonal in time, we included all motor metastates as regressors as well as a mean and linear trend features. A single design matrix per dance was constructed to generate one regression coefficient image per meta-state, per dance. Example design matrices can be found in the Supplementary Material.

### 2.6) Second-level analysis

#### Motor block design

In MEG studies it is typical to contrast the movement period to a baseline (rest) epoch within the same trial (e.g. Pakenham et al., 2020). We replicate this approach in our second-level GLMs. Each subject was tested individuality, such that the ‘scans’ were the individual trials for a single subject. For each limb, the 1st level regression coefficient images (either block design or HMM timed) were put into a paired-T test design matrix with their trial-specific rest regression coefficient images and a T-contrast between the active and rest images were generated, forming a paired-T image for each limb.

### Dancing

With no obvious hypothesis as to where a ‘rest’ period would occur in the data, we opted to contrast different movement states to each other (Ma et al., 2022).). Each scan here was an individual dance, meaning that instead of 5 subjects sent to the second level we had 20 dances instead (i.e. each dance (rather than each subject) was treated as an independent observation). Regression coefficient images from each dance were first put into a 1×5 factorial design matrix and T-contrasts between various limbs were generated.

For all T-images a whole brain Family Wise Error (FWE) correction using volumetric random field theory (Worsley et al., 1996) was applied using SPM.

## 3) Results

### Motor block design

*We* set out to assess whether in a controlled, block-designed motor paradigm we could detect participant movement in the video telemetry and use that to localise movement-related activity in the brain. Figure 2 depicts the results from a typical participant decoding the telemetry of the block-design experiment, where 1 of the 4 limbs were moved during a visual cue period. Similar results from the other participants can be found in the Supplementary Material. Figure 2A shows the binary timecourse for each condition of the experiment and the closest matching HMM state timecourse. Here the metric for matching a state to the original timecourse was the Jaccard index, with a mean±SD index for the selected 4 states of 0.65±0.06 (p < 0.001, see Supplementary Material). The resultant HMM states qualitatively resemble the original experimental timeseries they were compared against. The regression heatmaps in Figure 2B show which key-points on the body were implicated in each state. We observe that the parts of the body the regression highlighted correspond to the pre-defined experimental condition.

One feature that the telemetry identifies is the reaction time between a stimulus being presented and the participant executing and concluding the movement. Figure 2C illustrates this with an example trial (where the participant was asked to moved their left arm) in which the black dashed line representing the stimulus timing, and the blue solid line the corresponding the posterior probability of state being active. Here we observe that the participant takes (approximately) an extra second to cease movement after they were cued to stop. We found that across all trials the total time movement was executed for was 4.2 ± 0.9 (mean ± SD) s. Figure 2C also contains an additional period where the state being active is highly active at around t=10 s, which was within the inter-trial interval. To illustrate why the state is active, we have overlaid the associated telemetry timeseries from that state (made of the weighted sum of the keypoints highlighted in Figure 2B). We see that after the large motions during the trial, smaller motions are detected in the telemetry, which the HMM is sensitive to. These two examples within Figure 2C highlight how the telemetry data can reveal when the participant deviates from the experimental design.

We quantitatively compare the decoding of the experiment to the prescribed experimental timings in Figure 2D-E. These should be interpreted in the context of the variable reaction times of the participants. Figure 2D is the confusion/classification matrix comparing the ground truth of the experimental condition to the HMM derived Viterbi path. The Viterbi path is binarised for each labelled experimental state. Note for the ‘rest’ state, this represents the remaining 11 unlabelled states from the HMM. For each condition the predicted state was the most dominant, and we get an overall Fi score (non-weighted average of the Dice coefficients; Manning et al., 2008) of 0.814 (p < 0.001, see Supplementary Material). For the movement conditions, we see that a proportion of the movement states are predicted to be the rest condition, which would correspond to the reaction time for the participant to initiate the movement after the cue. Figure 2E shows the Receiver Operatic Characteristic (ROC) curves for the HMMs probabilistic state timeseries. Again, the timeseries for the unlabelled states are pooled together to represent the ‘rest’ condition. Here we are assuming a one-versus-all classification (i.e. correctly classifying a given state versus not). The areas under the curve (AUC) for each of the states are high (left leg, 0.868; right leg 0.862; left arm, 0.975; right arm, 0.990; rest, Q.8684) resulting in a macroaverage (non-weighted average) AUC 0.912 (p < 0.001, see Supplementary Material). We also performed a split-half cross-validation, which showed similar levels of performance (see Supplementary Material).

Figure 3 shows the T-contrasts where 8-30 Hz oscillatory power in rest periods were significantly higher (p < 0.05, FWE corrected) than the original experimental-derived active epochs (Fig. 3A) or the HMM-derived epochs for each of the 4 experimental conditions (Fig. 3B). For clarity, we have applied a threshold of 70% of the most extreme T-statistic in the image. In all conditions, we observe the characteristic event related desynchronisation (ERD) of power during movement. Focusing on the block-design derived results (Fig. 3A), we first observe that all of the peak locations are in the precentral gyrus (motor cortex). Second, we see that all 4 of the images follow the traditional organisation of the motor cortex (Gordon et al., 2023), with the arm condition peaks localised more lateral, and the leg peaks appearing medially. The peak T-value locations can be found in Table 2. Considering the results of the HMM-derived timings we note two things. First, the peak locations are very close between the two approaches (distances between peaks no further than 9.2 mm apart, see Supplementary Information). Second, the threshold statistical maps are similar in their morphology and their T-statistics are of a similar magnitude to the block design images. To summarise, the telemetry-based analysis gives comparable results to the block design.

**Figure 3.**
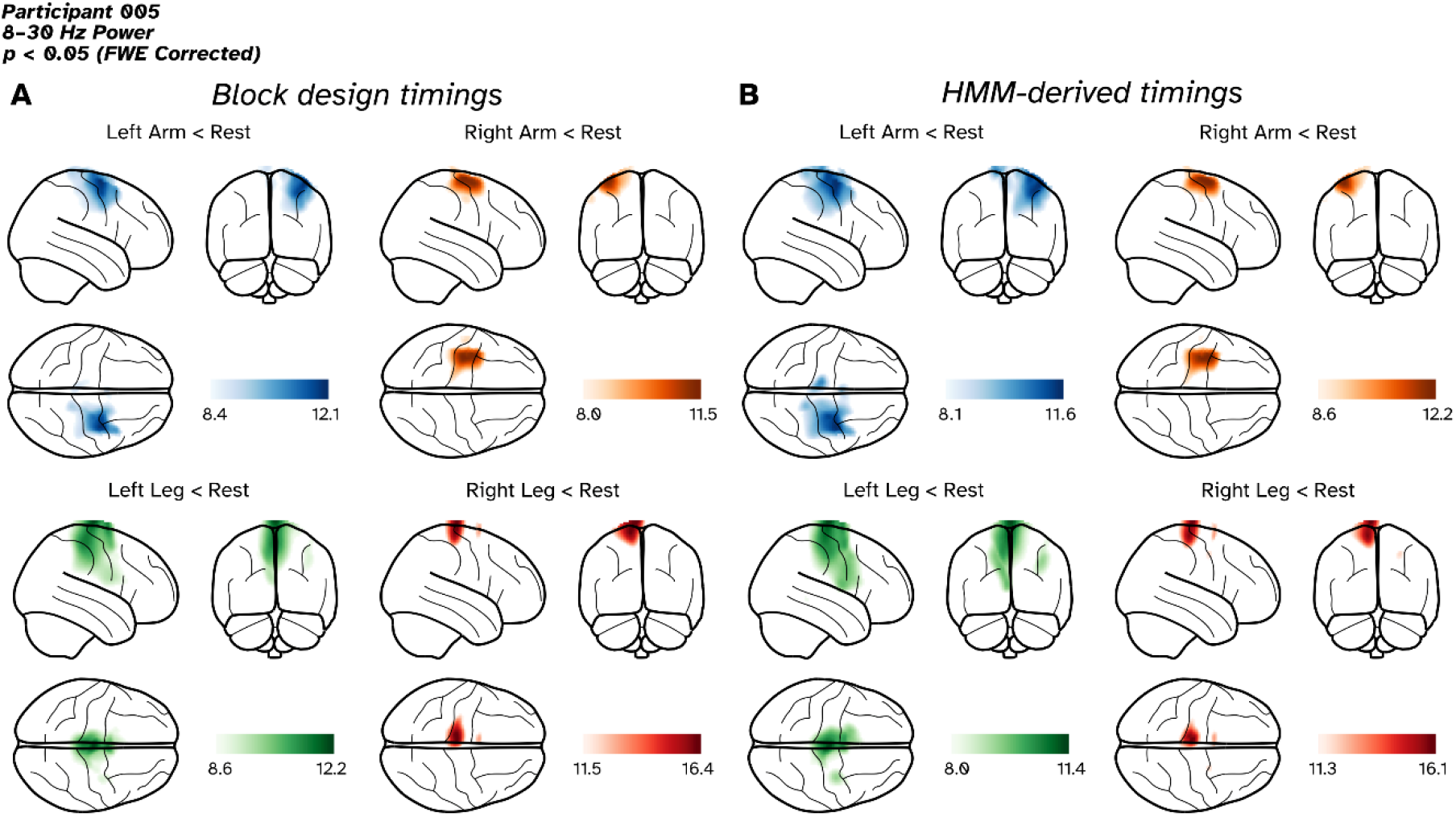
Source localisation results of a block designed, 4-condition motor experiment in a single subject, with results presented a glass brain. A) T-contrasts for the 4 active conditions v. rest based on the timings specified by the block design. B) T-contrasts for the 4 active conditions v. rest based on the HMM-derived timings. Images have been threshold to show anything within 70% of the most extreme statistic.

#### Dancing

Having established that our video telemetry pipeline works with a traditional, constrained motor paradigm, we extended our investigation to a series of more complex and naturalistic movements during a choreographed dance routine. Of the 25 states extracted from the telemetry, 13 were categorised into 5 meta-states, based on which key points on the body were employed. These meta-states were used to epoch the MEG data for further analyses. The heatmaps of these meta-states are depicted in Figure 4A. These represented each of the 4 individual limbs and an additional state representing both arms being moved in unison. Individual maps and timeseries of the 25 states can be found in the Supplementary Material. Figure 4B shows the fractional occupancy of a meta-state across the 15 sessions, which reveals a clear temporal structure emerging from the dance across all subjects and sessions. We see that each of the individual limb states dominate for a verse and chorus and crucially the dominant state corresponds to the theme of the verse *(“You put your left arm in”* for verse 1, right arm for verse 2 etc). We can also observe when a participant switches from one limb to another within a verse. For example, after 16 seconds where people transition from moving one arm to both at the line *“you do the hokey-cokey”* the dominant state switches from the left arm state (blue line) to the both arms state (purple line).

**Figure 4.**
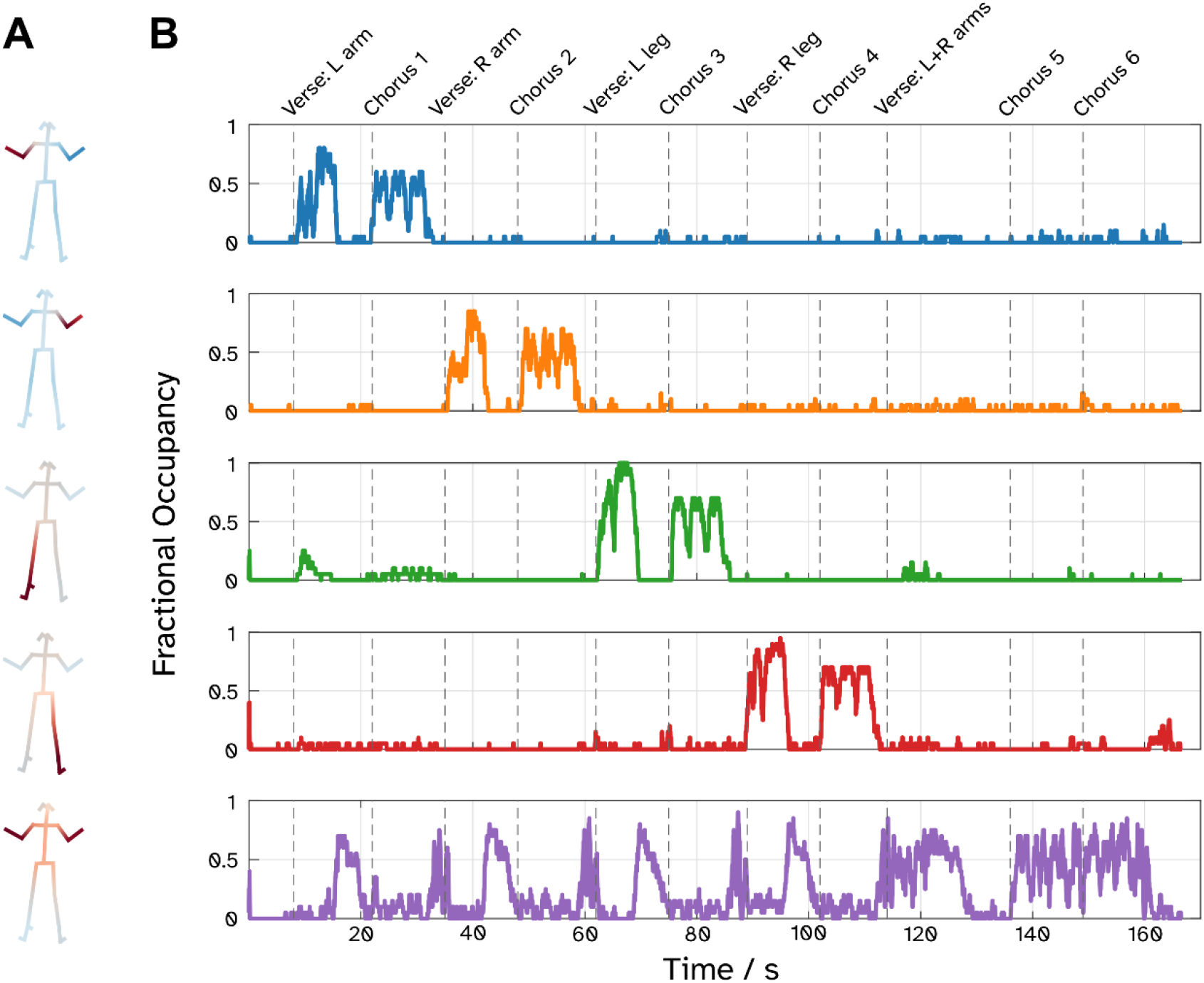
Five telemetry states extracted from the video of the 4 participants dancing to the Hokey Cokey. **A)** The heatmaps related to regressing the binarised state timecourses against the key-point velocity data, deep red areas show the parts of the body represented the most by each state. **B)** Session-averaged timecourses for each state, generating fractional occupancy timeseries, revealing the onset and offsets of dominant states over the progression of the dance.

Activation maps in the 8-30 Hz band, as defined by contrasting different movement states, are depicted in Figure 5. In particular we focus on three main contrasts of interest. For Figures 5A and 5B, we generated paired-T contrasts, where both arms moving (purple state in Figure 4B) were subtracted from one of the single arm states. First, we contrast when the *left arm* state had more power than the *both arms* state (Fig. 5A) and the resultant paired-T image shows significant activation (p < 0.05, FWE corrected) over the left dorsal sensorimotor areas (associated with right-upper limb movement). To get an activation of the right arm may be counterintuitive, but we note that contrasting both arms to a single arm should result in the activation map of the un-contrasted limb from a set theory perspective (the contrast represents the disjunctive union of the two limbs). Further supporting this idea is the opposite contrast of *both arms* to *right arm* (Fig. 5B), where we see a significant effect in the opposite hemisphere, representing left-arm movement. We note that we also contrasted single arm movements against each other, these are found in the Supplementary Material. Finally, we localise activity related to the movement of the legs in Figure 5C. We summed the *left leg* and *right leg* images to make a pooled mean effect of leg activation, finding significant activation in medial areas of the brain corresponding to lower limb movement. In summary, we have been able to extract movement timings from a complex motor paradigm and recover a plausible representation of motor activity in the brain.

**Figure 5.**
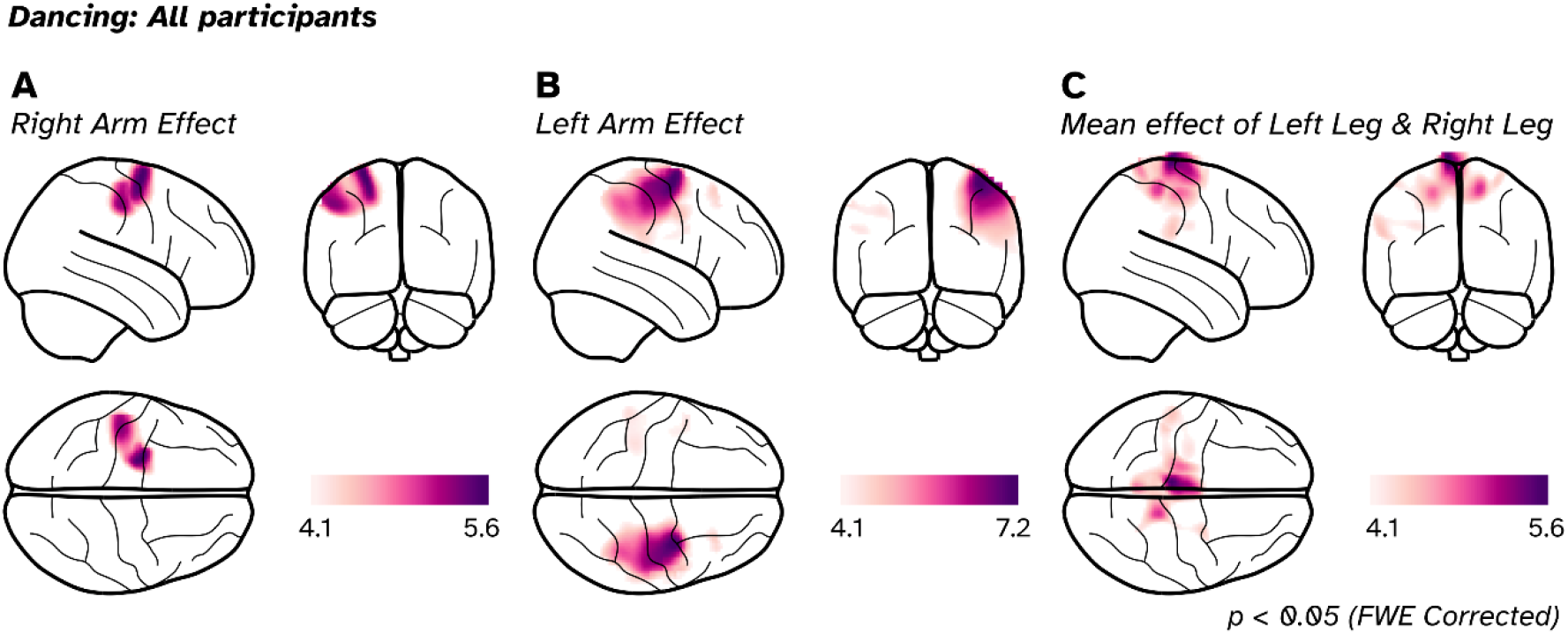
Group activation effects of 8-30 Hz power contrasted between different motor states from participants dancing, shown on glass brains. **A)** Right arm effect, derived by contrasting the left arm to both arms. **B)** Left arm effect, derived by contrasting right arm to both arms. **C)** Combining the left and right leg states contrast to get the mean effect of both legs reveals reductions in oscillatory power during movement over the medial motor areas. For all images the threshold was set to a T-value where p < 0.05 (FWE corrected).

## 5) Discussion

Wearable MEG systems (such as those using OPMs) are promising technologies for the future of naturalistic, mobile neuroimaging (Stangl et al., 2023). Work on making wearable MEG systems compatible with immersive virtual environments (Roberts et al., 2019) and ambulatory movement (Holmes et al., 2023b; Mellor et al., 2023; Seymour et al., 2021) is already well underway. However, the collection, analysis and fusion of complex behavioural data during the experiment with MEG data is something that has not yet been addressed. Here we have introduced a processing framework of movement data to demonstrate how this could be approached.

Combining motion capture data (either from motion capture suits or inertial measurements) has been combined with non-invasive functional neuroimaging regularly in the past (Lorenz et al., 2024; Mazurek et al., 2020; Miyakoshi et al., 2021), but due to the stationary nature of cryogenic MEG systems has been off-limits to MEG until OP-MEG became available (Hill et al., 2020; Seymour et al., 2021; West et al., 2024). However we gone one step further and removed the need for any additional hardware to be placed on the participant to track their motion. We have shown that it is possible to extract precise experimental timings to process OP-MEG data, entirely from marker-less decomposition of behavioural data, derived with open-source machine learning approaches applied to videos of participants executing limb movements. This the first time pose estimation has been combined with MEG and joins only a handful of other invasive and non-invasive functional imaging studies which have implemented such an approach (McDermott et al., 2023; Singh et al., 2021).

We first tested its capabilities in a well-controlled, block-design experiment and found it could recover the experimental timings needed to identify task-based changes in neural activity. We then applied this approach to a dancing paradigm, where after some basic choreography instruction to the participants, all movement was based on the interpretation and timing of each individual. From these video recordings, we derived neuronal states associated with different dance moves, which were mapped onto distinct representations of the limbs along the sensorimotor cortical surface. Our approach is particularly applicable for naturalistic studies, where consistent timings across participants are not guaranteed. Had we set experimental timings to the lyrical cues played to the participants, it would have overlooked the variability in subjects’ actual moves during the dance. By decoding these motor states from the behavioural data, we recover the subject-specific timings and factor in what could be previously described as ‘noise’ in our experimental design. This had been identified as one of the key challenges to developing successful mobile, naturalistic neuroimaging studies (Stangl et al., 2023).

Using regular (visual spectrum) video data to analyse movement with neural-network-based pose estimators (Bazarevsky et al., 2020; Cao et al., 2021; Mathis et al., 2018) can forgo the need for additional reflective markers on the body. This has the additional benefit for the family of OPMs we use, which use infra-red laser light to operate (and can be subject to interference from other light sources). The use of Hidden Markov Models (HMMs) to partition motion telemetry data is not uncommon (Agrahri et al., 2022; Buderman et al., 2021; Conners et al., 2021; Singh et al., 2021), as they are able to exploit the temporally-rich nature of the multivariate motion data. It should be noted that we need not have used HMM if a specific limb (eg left-hand) were the question, as most of the motion could be captured from a single keypoint. The advantage of HMM is that it allows us to express complex multi-variate changes of posture in a semi-supervised manor. This has potentially large implications for the clinic in (for example) telemetry recordings for epilepsy where many hours of video and encephalographic data can be collected. As potential alternatives to HMM, we note that note that principal or independent component analysis could also be ways forward.

Our results clustered into locations of the upper limb movements (dorsal motor cortex) and lower limb areas (medial motor cortex) that conform with the functional neuroanatomy of the sensorimotor cortices (Gordon et al., 2023; Jensen et al., 2022). This separation was clear in the block-design motor experiment, where the participant was seated and it was easier to move a limb in isolation. For the dancing paradigm, subjects were moving their whole bodies during each dance move, and never just moved one limb in isolation. This explains, for example, why there was activation within lower limb areas only when contrasting the leg states together rather than subtracting from each other, as muscles in both legs are activated when executing the swing movement with one leg. This investigation provides a proof-of-principle of the potential of wearable MEG to facilitate functional neuroimaging without highly controlled behavioural tasks.

As well as impacting naturalistic neuroscience research, approach also have clinical value, assessing pose-based behavioural data has been shown to be beneficial in assessing Parkinson’s disease (Kaneko et al., 2024; Roth et al., 2021; Sabo et al., 2022), the progress of Friedreich’s ataxia (Kadirvelu et al., 2023), as well as detecting nonverbal behaviour in Autistic Spectrum Disorder (Kojovic et al., 2021). Furthermore, combining pose-estimation with a flexible neuroimaging system such as OP-MEG promises to improve our understanding of these conditions in ways that were previously unavailable. There are direct clinical pathways where this could offer immediate benefit, such as in paediatric cases. For example, cases of refectory (drug resistant) epilepsy where surgical intervention may require careful localisation of the eloquent cortex with functional neuroimaging, to ensure that key faculties are not impaired when removing the seizure onset zone. This is typically assessed with functional MRI (Al-Arfaj et al., 2023; Barras et al., 2016), but success of the mapping is limited to the performance of the patient, and their tolerance of the MR system. For young children with epilepsy, this effect may be compounded (Yerys et al., 20G9). Solutions to this could include trying to map the motor areas in the resting state (Krishnamurthy et al., 2022), but there may be a more naturalistic alternative. Children playing when undergoing video-M/EEG telemetry sessions for seizure monitoring will enter states of movement and periods of rest naturally. If the video footage was processed to demarcate these periods it is entirely feasible that these motor maps (and indeed language or other key regions of eloquent cortex) could be derived *for free* out of long (often several hours) recordings where the primary goal is to locate ictal/interictal activity. Using play in neuroimaging may also be favourable for other populations, such as investigating the progress of motor development in children with Autistic Spectrum Disorder (An et al., 2021; Wilson et al., 2018), where a flexible imaging system which can adapt to the child, whilst simultaneously tracking movement should lead to more successful recording sessions than a large unadaptable system.

One technical aspect of the OP-MEG acquisition is the lack of additional active shielding applied to our OPM recordings. Our magnetically shielded room provides adequate shielding in the centre of the room to provide a low enough field to keep the sensors in their operational range (up to 4.5 nT from their initialisation point). However, we did not account for cross-axis projection errors (CAPE; Borna et al., 2022), which become apparent in recordings when the background field deviates by approximately 1 nT or more from the sensor’s zero point. Cross-axis projection errors result in additional asymmetrical non-linearities in the field-to-voltage response of the sensors (Schofield et al., 2023), which in turn has an impact on source localisation performance (Borna et al., 2022). In our block-designed motor experiment, the largest field experienced by any sensor was 2.1 nT from its zero-point. For the dancing data, the largest fields measured for each of the 5 subjects were 1.5 nT, 0.6 nT, 1.2 nT 2.6 nT and 3.5 nT from their zero-point. The data presented here will contain CAPE effects, with a worse case scenario of 3.5 nT generating approximately a 20 % gain error or in terms of source localisation, about 8 mm localisation error (estimates taken from Borna et al., 2022). However, we believe our results are not critically compromised by CAPE. First our environmental interference reduction method (HFC) is robust to the nonlinearities introduced by CAPE (Tierney et al., 2022), secondly the similarity of the results across subjects from the first task and between tasks suggests CAPE does not invalidate our findings. Methods which counteract field changes experienced by the sensors during ambulatory motion, whether through dedicated nulling coils built into the shielded room (Holmes et al., 2023b) or by using a dynamic closed-loop system built directly into the sensors (Lee et al., 2014; Mellor et al., 2023; Nardelli et al., 2019; Robinson et al., 2022) could be employed for future studies to control for CAPE-related nonlinearities.

One might query the wisdom of using multiple generations of QuSpin OPMs on our final participant. Both sensor types are designed to be compatible with each other. Furthermore, once calibrated, the noise floors are of similar levels, in fact the variability within generations is as high as between generations. This means we treat the third-generation sensors, once initialised essentially as a second-generation sensors with a bonus axis. The need for data whitening prior to analysis is not essential. Further other studies have also used this setup successfully (Corvilain et al., 2024; Spedden et al., 2024; West et al., 2024) so we believe that the combination of the sensors themselves is not a confound.

To conclude, our work provides additional support for the exciting opportunity of OP-MEG for studying the neural basis of complex motor functions, but also spatial navigation, memory and social interactions in realistic and ecologically valid situations, in both health and disease.

## 6) Acknowledgements

Funding for this project was provided by EPSRC (EP/T001046/1) from the Quantum Technology hub in sensing and timing (sub-award 0TPRF02) awarded to GRB and MJB. SM is funded by EPSRC-HIP award (EP/V047264/1). NAA, RAS are supported by a Wellcome Principal Research Fellowship (210567/Z/18/Z). TMT is funded by a fellowship from Epilepsy Research UK and Young Epilepsy (FY2101). MFH is supported by a Non-Clinical Post Doctorial Fellowship from the Guarantors of Brain. MS is supported by a Wellcome Technology development award (223736/Z/21/Z). GCO and DB are funded by a UKRI Frontier Research Grant (EP/X023060/1). The Wellcome Centre for Human Neuroimaging is supported by core funding from Wellcome (203147/Z/16/Z). The GPU was supported by an NVIDIA academic hardware grant awarded to MFH.

## 8) Data and code availability

The MATLAB scripts used to analyse the data are available at https://qithub.com/qeorqeoneill/videomeq. Openpose 1.7.Q is available to download from https://qithub.com/CMU-Perceptual-Computinq-Lab/openpose. The data, downsampled to 2 kHz (with antialiasing applied) is available to download from Zenodo at https://doi.org/1Q.5281/zenodo.8139849. Openpose telemetry data from the video is included. To conform with data protection regulations, we cannot share the original video on which the pose inference was performed.

## 9) Contributions

**GCO:** Conceptualization, Methodology, Software, Formal analysis, Investigation, Writing -original draft. **RAS:** Methodology, Software, Resources, Writing - Review and Editing. **SM:** Software, Resources, Writing - Review and Editing. **NA:** Software, Resources, Writing - Review and Editing. **TMT:** Software, Resources, Writing -Review and Editing. **LB:** Investigation, Resources, Writing -Review and Editing. **MHF:** Resources, Writing -Review and Editing. **MES:** Investigation, Resources, Writing -Review and Editing. **RCT:** Software, Investigation, Resources, Writing -Review and Editing. **DB:** Resources, Writing - Review and Editing. **SB:** Conceptualization, Methodology, Resources, Writing - Review and Editing. **MJB:** Conceptualization, Writing - Review and Editing, Funding Acquisition. **GRB:** Conceptualization, Methodology, Software, Resources, Writing - review & editing, Funding acquisition.

