## Supplementary Material for "Combining video telemetry and wearable MEG for naturalistic imaging"

George C. O'Neill[a,b], Robert A. Seymour[b], Stephanie Mellor[b], Nicholas Alexander[b], Tim M. Tierney[b], Léa Bernachot[b], Mansoureh Fahimi Hnazaee[b], Meaghan E. Spedden[b], Ryan C. Timms[b], Daniel Bush[a], Sven Bestmann[b,c], Matthew J. Brookes[d], Gareth R. Barnes[b]

[a]*Department of Neuroscience, Physiology and Pharmacology, University College London, London, UK*
[b]*Department of Imaging Neuroscience, UCL Queen Square Institute of Neurology, University College London, London, UK*
[c]*Department of Clinical and Movement Neurosciences, UCL Queen Square Institute of Neurology, University College London, London, UK*
[d]*Sir Peter Mansfield Imaging Centre, School of Physics and Astronomy, University of Nottingham, Nottingham, UK*

---

#### *1) Sensor layouts*

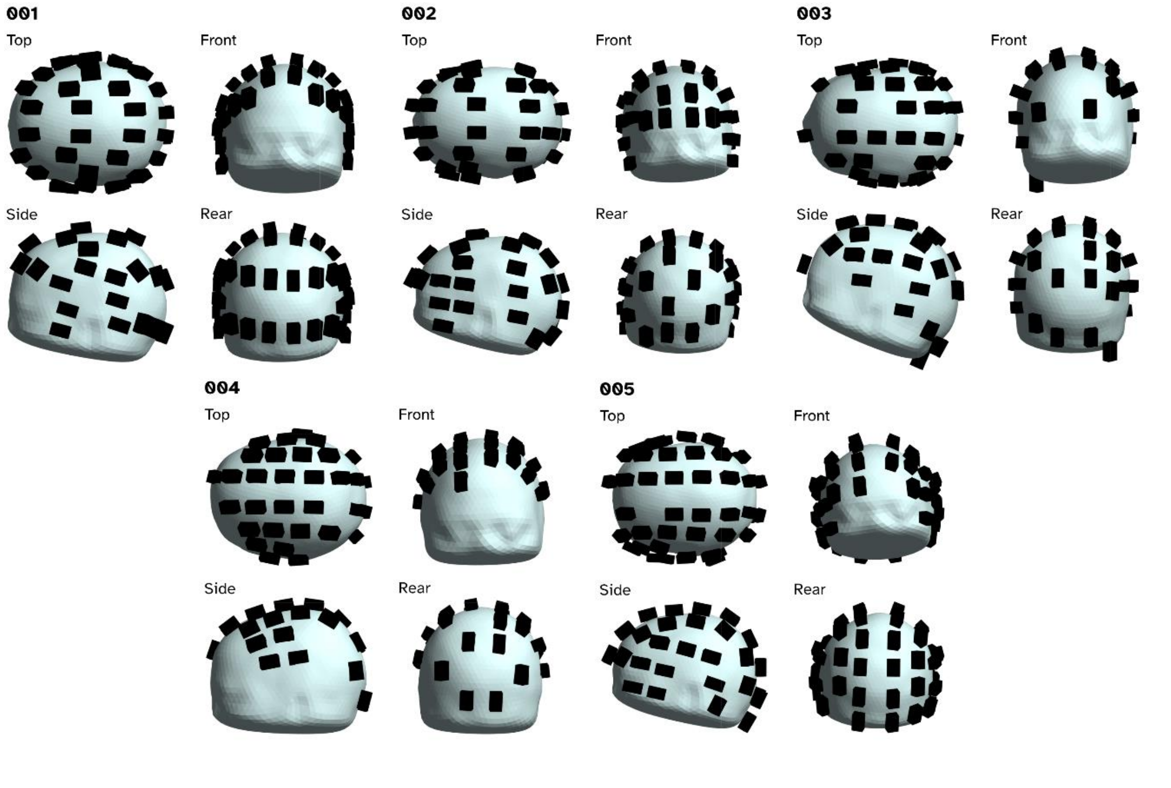


**Figure S1** - Renders of the sensor locations for the 5 subjects involved in the study, black cuboids represent the casings of the QuSpin magentometers, whilst the light blue meshes represent the scalps of the participants.

| Participant | Motor experiment | Dancing experiment | No. dances | QZFM Gen-2 | QZFM Gen-3 | Total Channels |
|---|---|---|---|---|---|---|
| 001 | N | Y | 3 | 43 | 0 | 86 |
| 002 | N | Y | 4 | 40 | 0 | 80 |
| 003 | Y | Y | 3 | 38 | 0 | 76 |
| 004 | Y | Y | 5 | 32 | 0 | 64 |
| 005 | Y | Y | 5 | 25 | 26 | 128 |

***Table S1*** *- Number of operational sensors and channels placed on the scalp for each participant. Each sensor provides one channel of axial-oriented field detection one tangential recording for 2$^{nd}$ generation sensors and a full vector measurement for the 3$^{rd}$ generation sensors.*

### 2) 1st level analysis design matrices

Figure S2 shows examples of the design matrices used in the study. The three left-most matrices represent those used for the motor block design experiment for an example trial. Here the key difference is how the first column is defined, either based on the timings of the block design (see Block Active and Block Rest) or via the HMM-derived timings (HMM Active). The rightmost column represents the design matrix for an example session of the dancing paradigm, where the first 5 columns represent the HMM-derived timings of the motor telemetry.

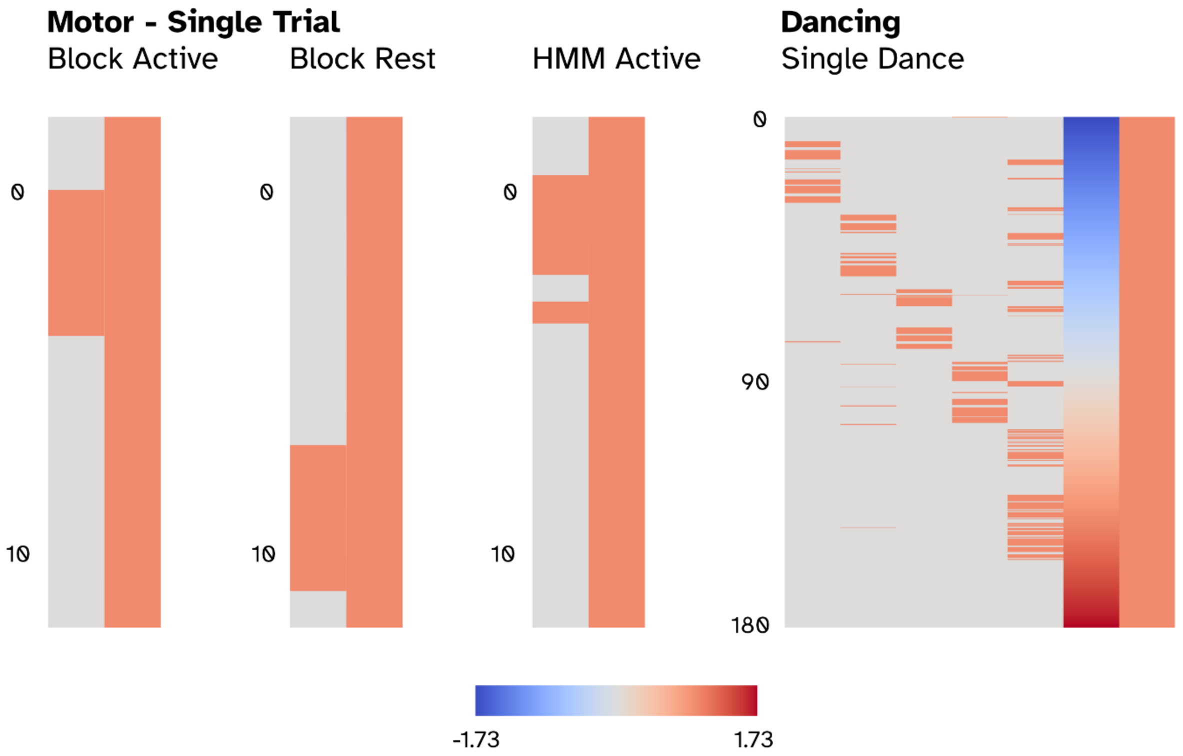


**Figure S2** – Example design matrices from the 1st-level analyses performed.

#### 3) ***Randomisation testing of classification metrics***

In the main manuscript we provided p-values for the Jaccard indices, $F_1$ scores and Area under curve (AUC) scores when classifying hidden Markov model (HMM) states to experimental conditions. Typically these scores do not have associated p-values for null hypothesis testing, so we developed a randomisation test to build a null distribution that each metric's score could be explained by pseudo-data.

Having fitted a HMM to the original telemetry data, we tested against a null hypothesis that random data which obeyed the fitted HMMs dynamics and distributions could explain the classification scores just as well. For a given iteration we generated a synthetic dataset of the same size where each state followed the multivariate Gaussian distributions fitted to the original data and state transitions were based on the fitted transition matrix. We then measured the Jaccard indices, F1 classification and AUC the synthetic dataset for the states which were labelled as the four limbs and rest in the original model compared to the real experiment. 1001 iterations of this were performed to build the null distributions below in Figure S3 for Subject 005. The three distributions show the sham results for the Jaccard indices (blue), F1 score (red) and AUC (yellow). Solid lines in blue, red and yellow represent the scores generated from the real data using the same HMM. All results exceed a $p < 0.001$.

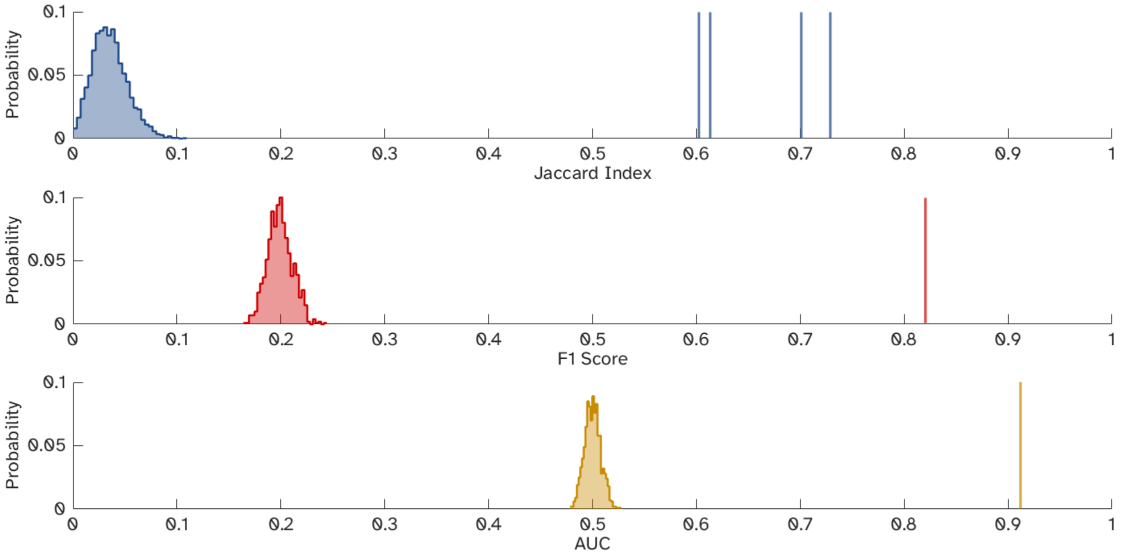


**Figure S3** – Results of the null permutation testing using synthetic data generated from the fitted hidden Markov model for Subject 005. The blue distribution represents the sham Jaccard indices, the red the sham F1 scores and the yellow the sham AUC results. Solid vertical lines represent the true scores from the participants data.

#### 4) ***Additional Motor Proof of Principle results from subjects 003 and 004***

Figures S4 and S5 are the results of processing the pose estimation telemetry from subjects 003 and 004 respectively. For each subject we present a pair confusion matrices (Figures S4C and S5C) and receiver operation curve plots (Figures S4D and S5D). The left hand plots are the metrics calculated between the block design and HMM output as per the main manuscript, whereas the right hand plots include a "reaction time correction". To explain our motivation we show an example set of trials from Subject 003 where the cue was moving their left arm in three consecutive trials (Figure S4). The black dashed line representing the cue to start/stop moving precedes when the HMM state for the right arm is active by a fairly consistent offset. Whilst the origins of the reaction delay (human or HMM) are not clear, we can quantify the delay (assuming the delay is consistent across all trials) using a cross correlation measure between the block design timeseries and the HMM Viterbi path. For Subject 003 this delay equates to 0.57 s and for Subject 004 it was 1.07 s. By shifting the block design timings by that those amounts we saw improvements to both the F1 scores in the confusion matrices and the AUC for the

ROCs. We also investigated whether Subject 005 had a systematic delayed reaction, whilst their reaction time were on average 0.33 s, there was no change the F1 or AUC scores by accounting for this, suggesting that their delayed onsets and offsets were more stochastic than systematic.

Figure S6-7 show the source location results for subjects 003 and 004, the resultant maps follow the same organisation of the sources we'd expect from sensorimotor areas (arms more dorsal, legs more medial) though we note that the results of subject 004 localise to the sensorimotor cortex rather than the motor cortex (which subjects 003's and 005's results do). All visualises sources exceed a $p<0.05$ (FWE corrected) significance threshold (which for a paired-T design with 30 trials, dof=29, critical T=4.33) The peak locations for each contrast (block design and HMM derived) can be found in Table S2

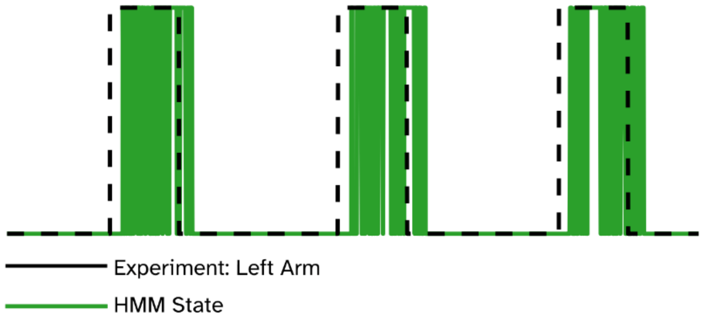


**Figure S3** – Example trials from Subject 003, where three left arm trials were presented consecutively. We see that between the cued experimental timings (black dashed line) and participant movement as defined by the HMM (green solid line) that there's a systematic delay of the onset and offset of subject movement.

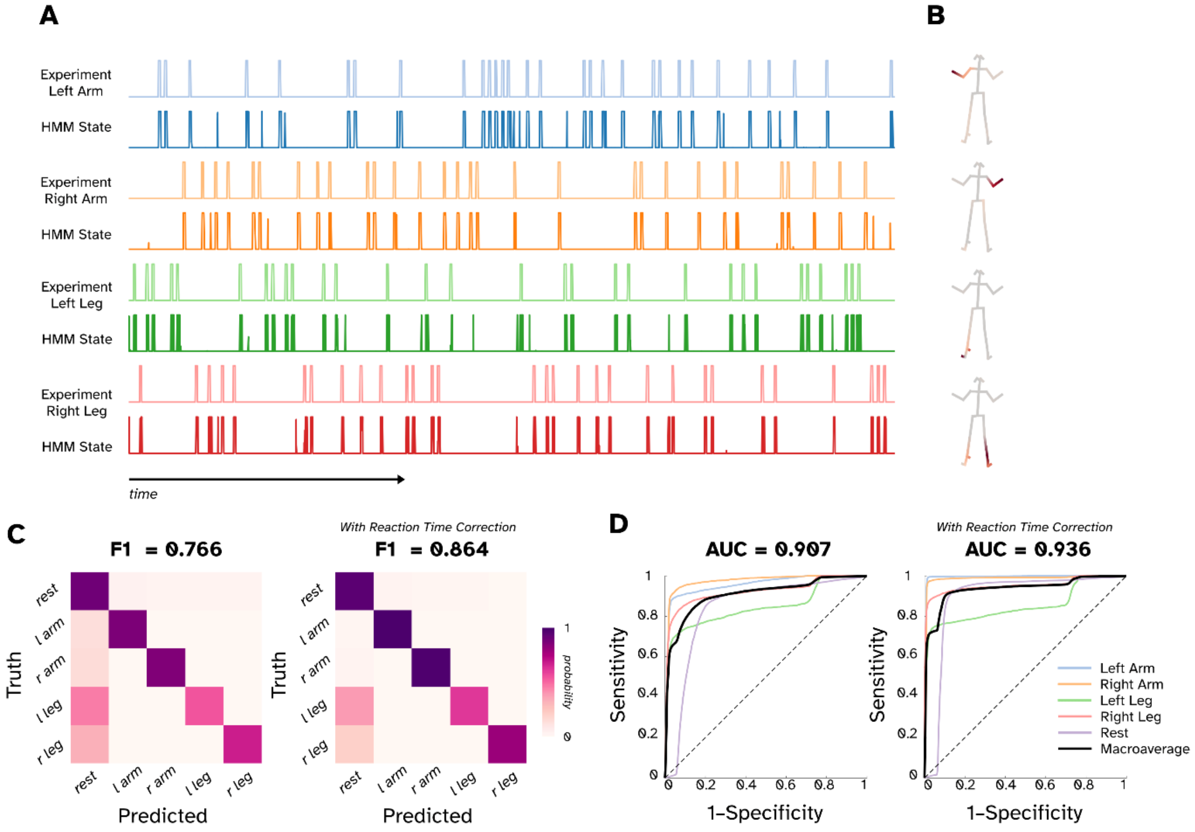


**Figure S4** – HMM telemetry results from Subject 003. A) The experimental block design with the corresponding HMM state probability timecourse. B) Regression heatmaps showing which keypoint locations are employed in a given HMM state. C) Confusion matrices showing the probability of the experimental block design and the HMM state overlapping. F1 classification score is improved by taking account of the systemic delay in reacting of the participant (right plot) compared to no correction (left plot). Receiver operation characteristics plots for both accounting for reaction time (right) and not (left).

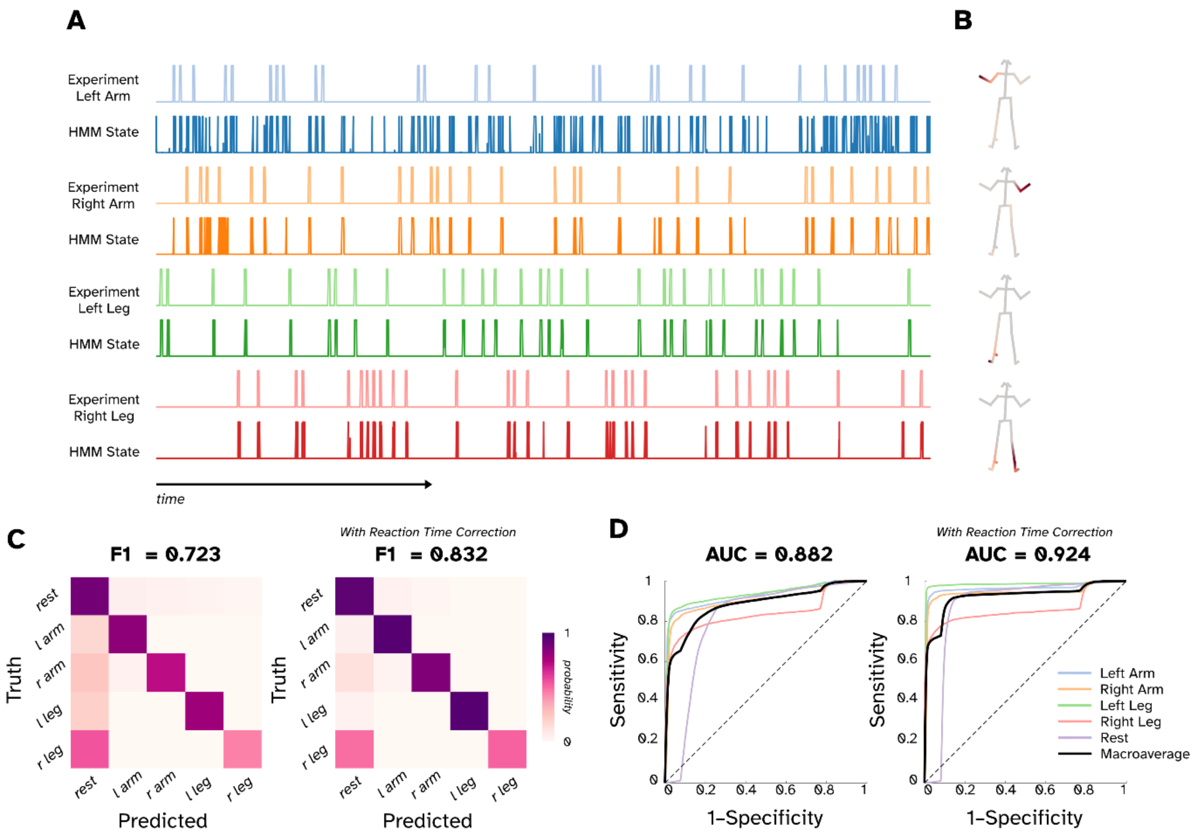


**Figure S5** – HMM telemetry results from Subject 004. A) The experimental block design with the corresponding HMM state probability timecourse. B) Regression heatmaps showing which keypoint locations are employed in a given HMM state. C) Confusion matrices showing the probability of the experimental block design and the HMM state overlapping. F1 classification score is improved by taking account of the systemic delay in reacting of the participant (right plot) compared to no correction (left plot). Receiver operation characteristics plots for both accounting for reaction time (right) and not (left).

***Subject 003***
***8-30 Hz Power***
***$p < 0.05$ (FWE Corrected)***

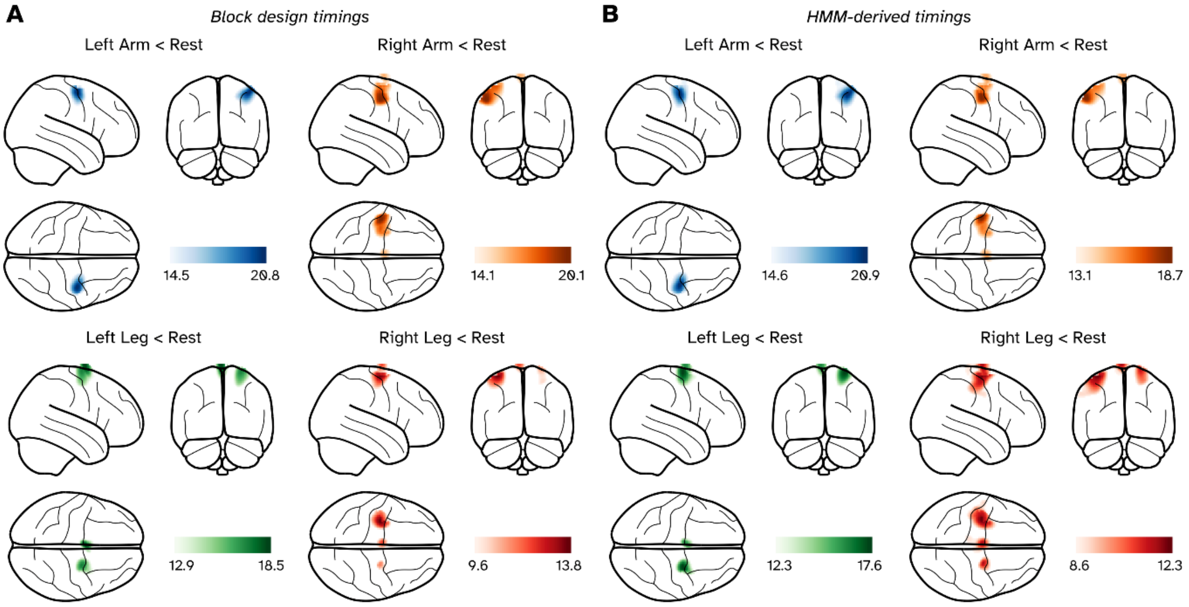


***Figure S6*** *- Source localisation results of the motor experiment in a subject 003, with results presented a glass brain. A) T-contrasts for the 4 active conditions v. rest based on the timings specified by the block design. B) T-contrasts for the 4 active conditions v. rest based on the HMM-derived timings. Images have been threshold to show anything within 70% of the most extreme statistic.*

***Subject 004***
***8-30 Hz Power***
***$p < 0.05$ (FWE Corrected)***

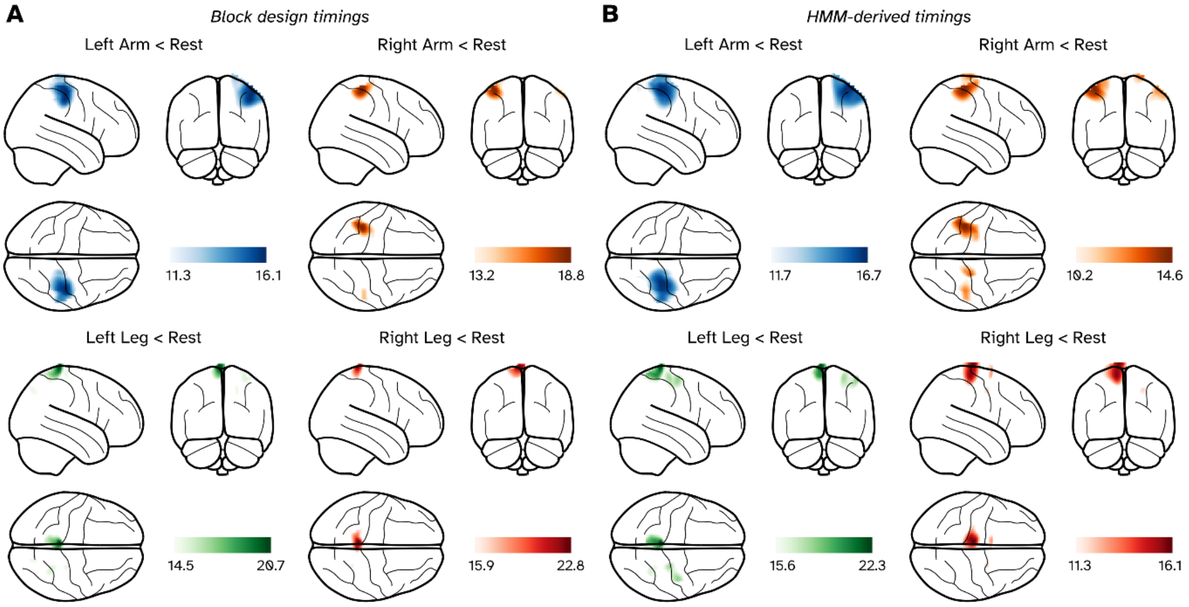


***Figure S7*** *- Source localisation results of the motor experiment in a subject 004, with results presented a glass brain. A) T-contrasts for the 4 active conditions v. rest based on the timings specified by the block design. B) T-contrasts for the 4 active conditions v. rest based on the HMM-derived timings. Images have been threshold to show anything within 70% of the most extreme statistic.*

| Participant | MNI peaks (mm) | Left Arm | Right Arm | Left Leg | Right Leg |
|---|---|---|---|---|---|
| **003** | Block design | 42 -8 58 | -50 -10 48 | -2 -4 80 | -34 -10 66 |
| | HMM | 38 -6 54 | -52 -10 50 | 28 -8 74 | -36 -10 66 |
| | Distance | 6 | 2.8 | 30.9 | 2 |
| **004** | Block design | 44 -24 54 | -38 -36 60 | -4 -40 80 | -6 -40 80 |
| | HMM | 36 -30 60 | -40 -34 62 | -4 -40 80 | -8 -38 80 |
| | Distance | 11.7 | 3.5 | 0 | 2.8 |
| **005** | Block design | 38 -4 66 | -38 -10 66 | 0 -18 82 | -12 -24 74 |
| | HMM | 38 -4 64 | -34 -6 66 | 0 -18 82 | -8 -22 82 |
| | Distance | 2 | 5.7 | 0 | 9.2 |

***Table S2*** *– Locations of peak activations for participants 003-005 using both the block design and HMM approaches.*

#### *5) Cross validating the Hidden Markov Model (HMM) on video telemetry*

In the main manuscript, we generated a confusion matrix to and ROC plot to quantify how well an HMM performed in decoding the original experimental conditions from the video telemetry from Subject 005. Whilst the data given to the HMM was unlabelled and it naturally partitioned the data into states which fit around the experiment, we understand quantifying the decoding performance on the data it was trained on is not good practice from a machine learning perspective.

To this end we performed a split-half analysis on the data. The trials in original block design experiment, whilst pseudorandomised were balanced to have equal loadings (15 of each condition) in each of the two runs, allowing us to fit models on one run and compare decoding performance on both runs. Figure S8A shows the ROC curves for each permutation of HMM, and we see that for the case where the HMM trained on run 1 decodes run 2 is the only case where the AUC is worse than the benchmark AUC in the main results (0.894, down from 0.912). In the cases where the models decode their own runs, we see marginal improvements to the AUC, though this result is unsurprising. The most interesting result of the HMM trained on run 2 used to decode run 1 gives the best performance, and AUC of 0.938, better than the model decode the dataset it was trained on. However, the differences in AUC are marginal. We see a similar trend with the confusion matrices' F1 scores in Figure S8B. In short, for this experiment the HMM generalises well enough to not have compromised decoding performance of the behavioural condition.

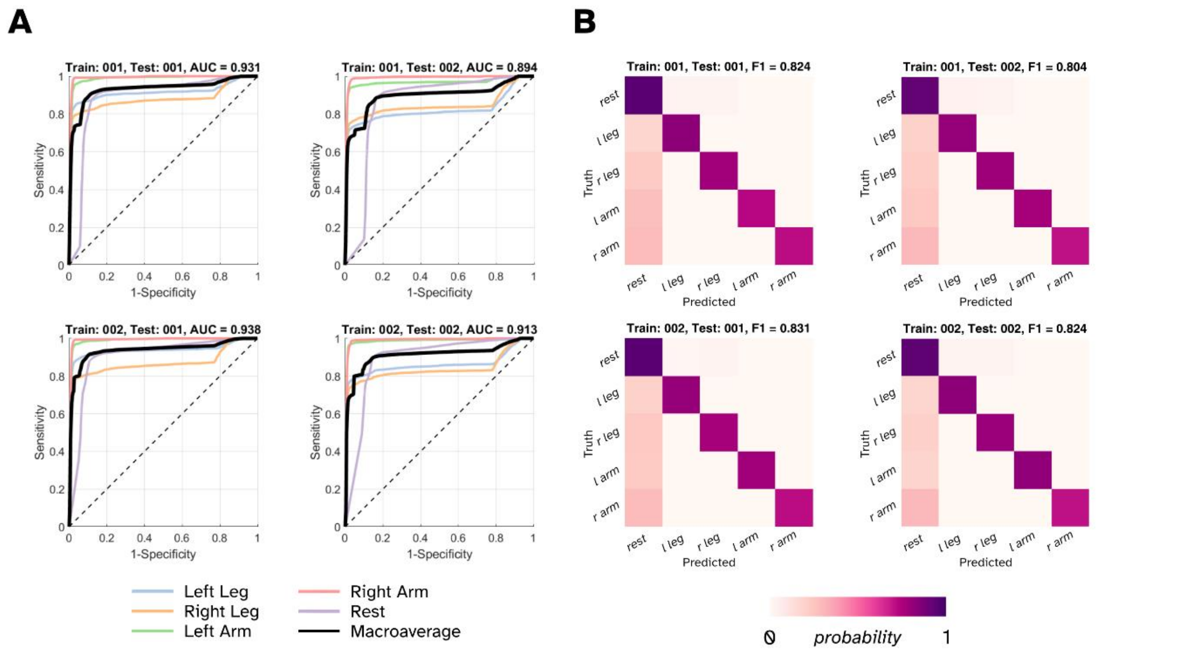


***Figure S8** - Results of classification performance of the HMM states in a split-half analysis. A) ROC curves showing the specificity and sensitivity of the states' probabilistic timeseries in a one v all classification. B) Normalised confusion matrices from the HMM Viterbi paths.*

### 6) *Additional contrasts from the dancing paradigm*

Figure S9 contains four additional which are referred to in the main manuscript. For clarity we've masked to the sensorimotor areas, something which the contrasts in the main manuscript did not require. Figures S9A-B show the contrast of the individual legs to the both arms state, whilst we do get a localisation which is medial motor areas and in the correct hemispheres, the T-values are too low to be considered to be significant. Figures S9C-D show the direct contrasts between arms and legs, here we've also visualised the negative T-values. Contrast one arm against the other (Fig. S9C) shows the particular "imbalance" in power changes for each arm. Whilst the cluster in the right hemisphere would be significant (critical T-value for $p < 0.05$ (FWE corrected) is 4.11), the cluster in the left hemisphere corresponding to the other arm is below this threshold (peak absolute T-value is 2.55). Likewise contrasting the legs to each other (Fig. S9D) shows lateralised asymmetry but no clear or significant interaction.

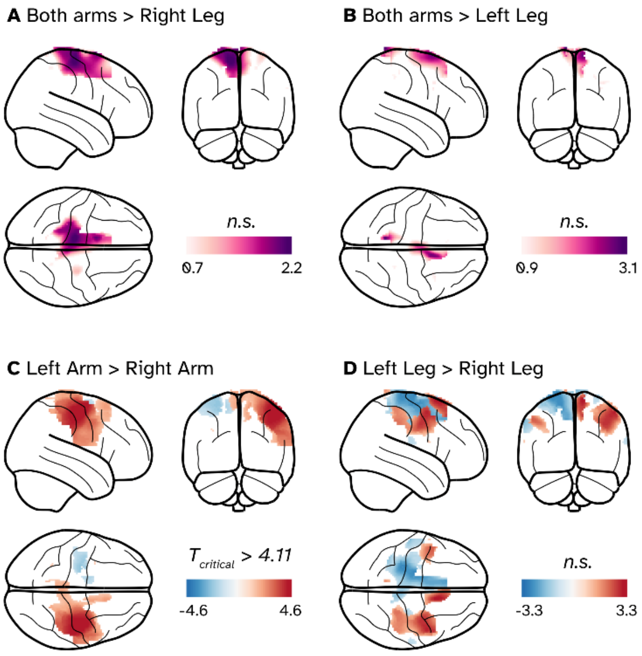


**Figure S9** – Additional T-contrasts from the dancing data.

### *7) HMM states from dancing*

Figures S9-S11 are the individual states generated from the group-concatenated telemetry from the dancing paradigm, where they have been grouped together by which meta-state they were ultimately labelled with. The left plot is the regression heatmap of the HMM Viterbi path regressed against the velocity of each key point; here the red represents and positive covariance with the HMM state, whilst blue a negative covariance. The right-hand plots are the trial-averaged probability of that state occurring across all dances.

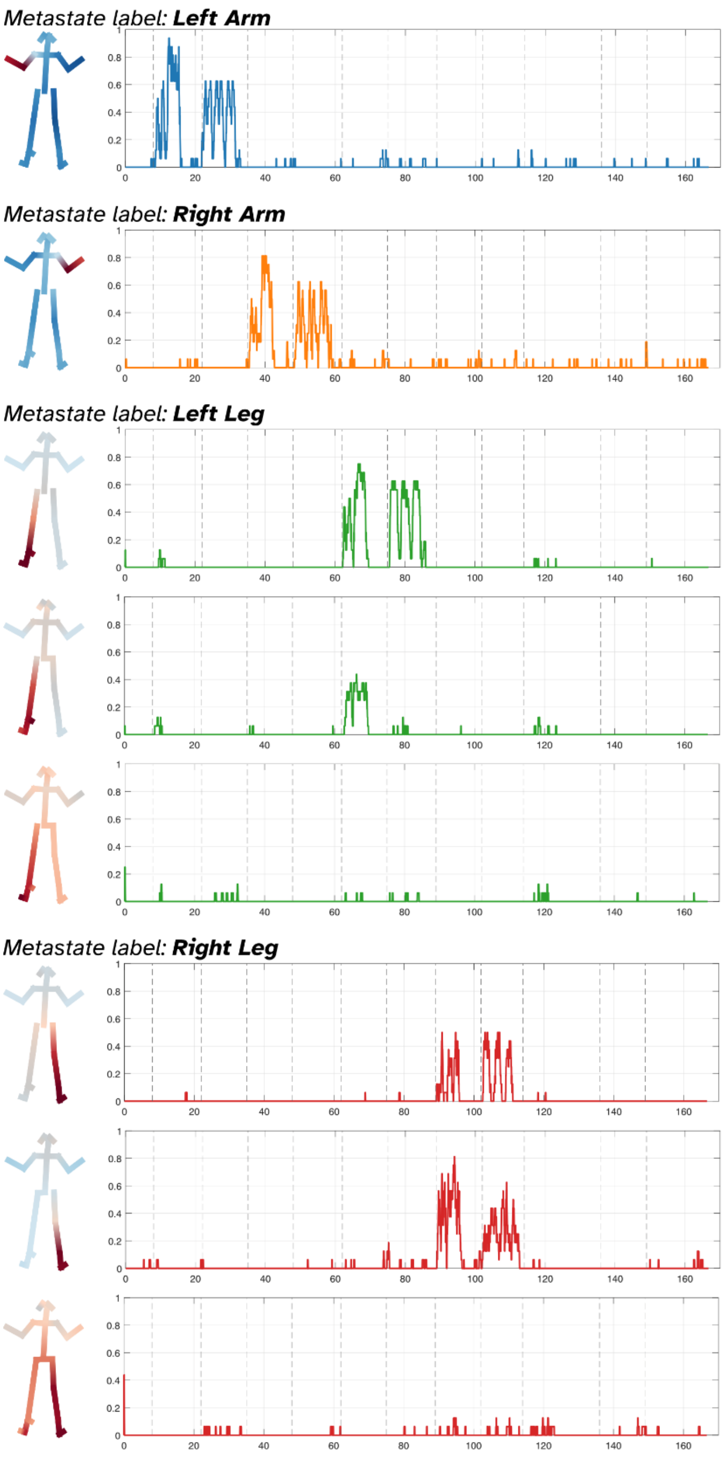


**Figure S10**

*Metastate label:* ***Right Leg*** *(cont)*

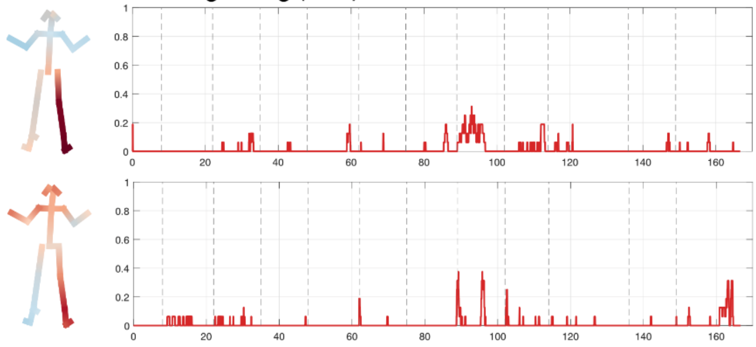


*Metastate label:* ***Both Arms***

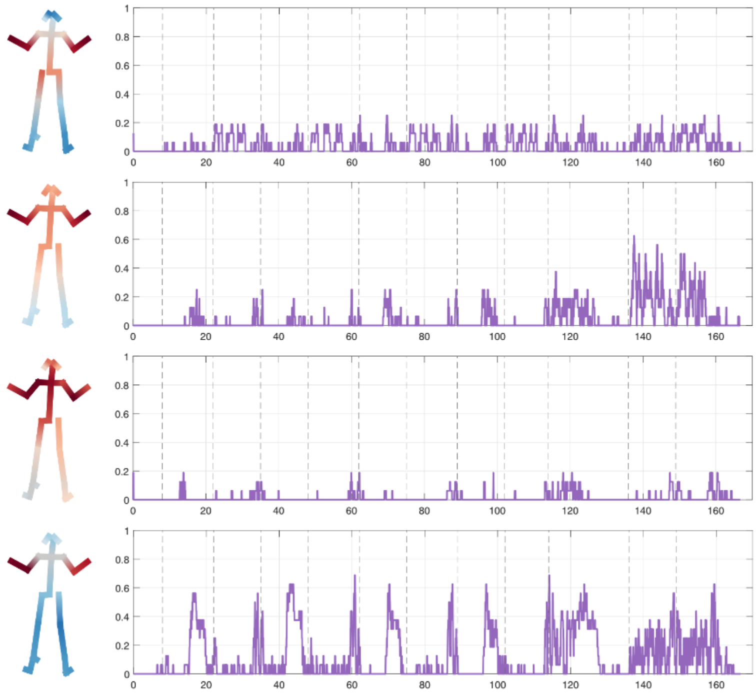


*Metastate label:* ***Unlabelled***

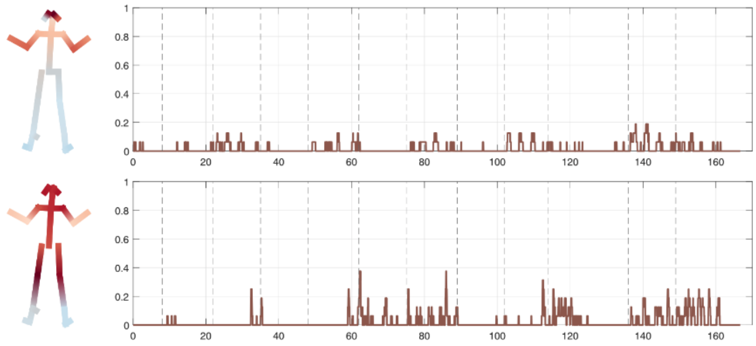


**Figure S11**

*Metastate label:* ***Unlabelled*** *(cont)*

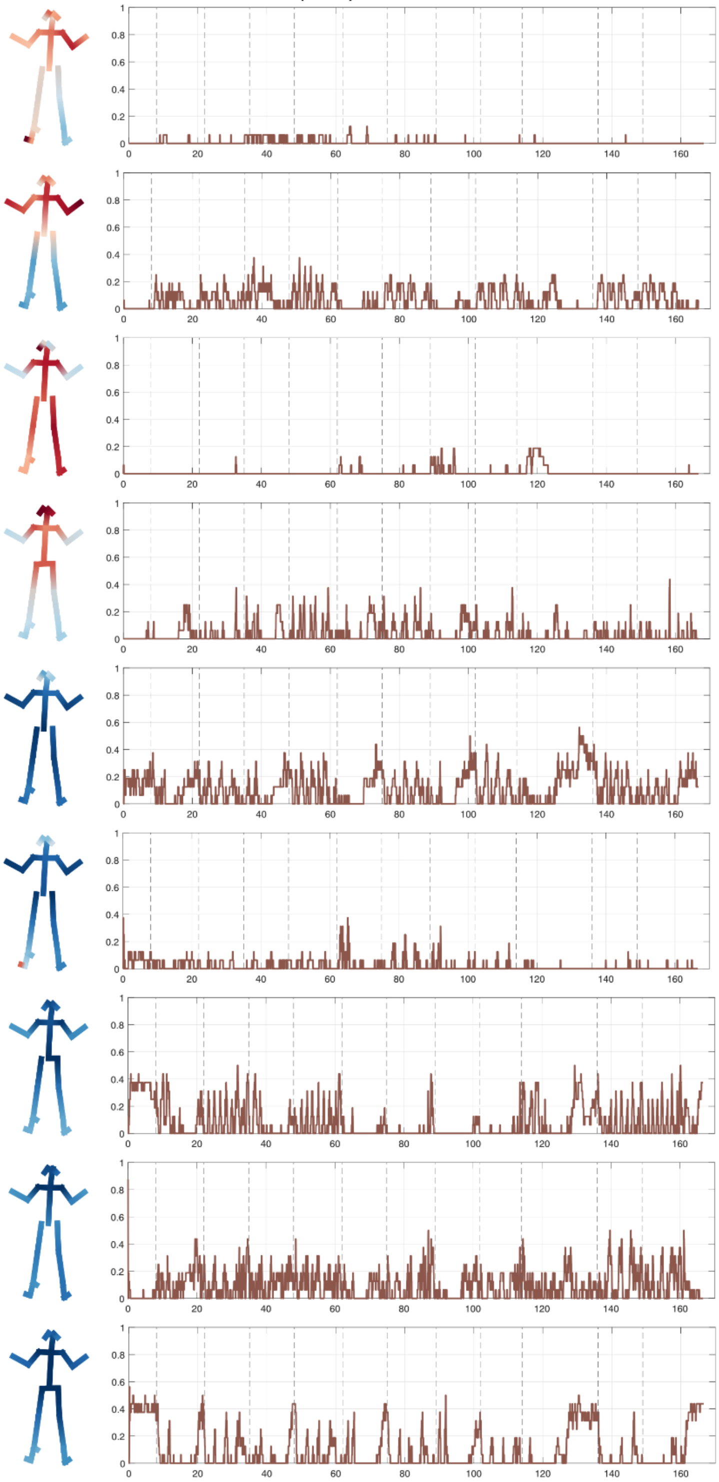


**Figure S12**
